# Detecting Large Indels Using Optical Map Data

**DOI:** 10.1101/382986

**Authors:** Xian Fan, Jie Xu, Luay Nakhleh

**Affiliations:** Rice University, Houston, TX 77005, USA; MD Anderson Cancer Center, Houston, TX 77030, USA; The Pennsylvania State University, Hershey, PA 17033, USA

## Abstract

Optical Maps (OM) provide reads that are very long, and thus can be used to detect large indels not detectable by the shorter reads provided by sequence-based technologies such as Illumina and PacBio. Two existing tools for detecting large indels from OM data are BioNano Solve and OMSV. However, these two tools may miss indels with weak signals. We propose a local-assembly based approach, OMIndel, to detect large indels with OM data. The results of applying OMIndel to empirical data demonstrate that it is able to detect indels with weak signal. Furthermore, compared with the other two OM-based methods, OMIndel has a lower false discovery rate. We also investigated the indels that can only be detected by OM but not Illumina, PacBio or 10X, and we found that they mostly fall into two categories: complex events or indels on repetitive regions. This implies that adding the OM data to sequence-based technologies can provide significant progress towards a more complete characterization of structural variants (SVs). The algorithm has been implemented in Perl and is publicly available on *https://bitbucket.org/xianfan/optmethod*.

## 1 Introduction

Structural variant (SV) detection is essential in understanding human genetic diseases such as cancer [11,37,23]. Detecting SVs is very challenging due to several factors, including the simple sequence context of the SV breakpoints [1], the multiple SVs aggregated to form a complex SV [39,31,2], and the repetitive nature of the human genome [13,34]. Advances in sequencing technology make it possible to detect SVs through computational tools [25]. Several SV detection methods using Illumina paired-end reads have been devised [6,16,41,18,32,8]. However, due to their small length (typically, 300bp read length), the focus was mainly on small indel detection and medium-sized simple SVs such as deletion and translocation [1,40]. Large SVs whose breakpoints fall at repetitive regions were not fully resolved by Illumina reads. PacBio single molecule reads [7,30], on the other hand, tackle those SVs in larger repetitive regions, and detectable SV types naturally generalize to insertion and inversion, due primarily to PacBio’s larger read length (typically 12kbp for RS II) [4].Nevertheless, the read length is still not enough for spanning large repeats, leading to missing SVs.

Optical Maps [33,21] produce one of the longest read lengths among all. It utilizes restriction enzymes to make fluorescent labels on the molecule wherever there is a 6 or 7bp sequence motif [17,3]. The molecule is then linearized and imaged. The subsequent image processing step measures the distance of the two neighboring fluorescent labels and outputs an array of integers, indicating the position (in bp) of each fluorescent label on the read. When the DNA has a structural variant with respect to the reference, the read has discordant patterns of integers with that of the *in silico* digested reference sequence. Read length is typically > 150kbp [24], which is one order of magnitude longer than PacBio reads and two orders of magnitude longer than Illumina reads. With such large length, Optical Maps data enables the detection of SVs that are missed by other technologies, and they have been applied to both normal and cancer patient samples [24,15]. Despite the large read length, computational methods are required for it to be widely used for SV detection by accounting carefully for OM data shortcomings, which include the small number of fluorescent labels in each read, and the various errors of additional labels (17%), missing labels (10%), and sizing difference [3].

The use of OM reads data for SV detection started from correcting [28], assessing [14] and scaffolding *de novo* whole genome assembly (WGA) from other sequencing technologies such as PacBio [30], Illumina [38], 10X [27] or a combination of multiple technologies [36]. Recently, there have been efforts for using OM alone for SV detection. There are two existing approaches to SV detection using OM data alone: assembly-based and alignment-based. In assembly-based methods, OM reads are assembled *de novo* into contigs, which are then compared with the *in silico* digested reference sequence [3,24]. Such de novo WGA strategy takes advantage of the randomness of errors in a cohort of reads for obtaining accurate and long contigs. However, due to the repetitive regions in the genome and the low resolution of the label coordinates, *de novo* WGA requires typically 70x read coverage in a diploid healthy human genome [12]. This makes it impossible to tackle those SVs with low coverage of reads. BioNano Solve [12] is an assembly-based approach and its recall is limited in low-coverage loci. Alignment-based methods, on the other hand, align the OM reads to the reference, and cluster the reads on focal regions where discordant patterns occur. OMSV [22] is an alignment-based method which uses the reads that can span the indels to infer insertions and deletions. It is computationally efficient as compared with BioNano Solve (at least one order of magnitude less time and much smaller required memory) and is applicable to loci with lower coverage of reads. However, in inferring indels, it can only cluster the reads having the same indel boundary. The design of OMSV limits the detection power only to indels in which a significant number of reads are confidently well aligned, but cannot deal with the indels when the aligners render different boundaries for different reads because of data noise (illustrated in Fig. 1). Mak *et al.* [24] combined the assembly-based and alignment-based approaches and used both WGA and an alignment-based approach for SV detection. However, their SV calling process involves heavy manual curation based on Illumina reads. Furthermore, no accompanying tool was released with the paper, making it hard to compare to other methods in a performance study.

**Fig. 1:**
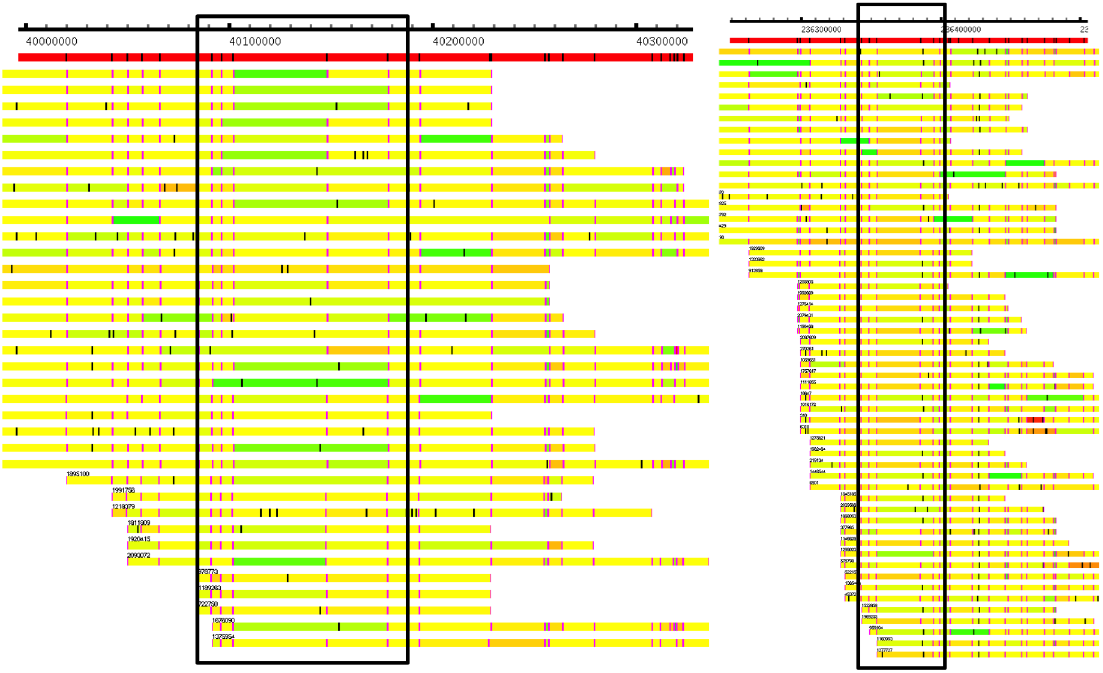
OMView [19] illustrations of a deletion (left), as validated by parents’ signal, and insertion (right), as validated by both parents’ signal and orthogonal sequence-based methods (i.e., the methods that are applied to sequencing technologies other than OM), both of which are missed by OMSV and BionanoSolve on NA12878. Shown is the alignment of the OM reads to the reference. Reference is the top bar in red. OM reads are the bars below. On the reference and OM reads, the vertical lines indicate the presence of a restriction enzyme fluorescent label. On a read, we call the part in between two neighboring restriction enzymes a fragment. Fragment length is the distance between these two restriction enzymes. A read is composed of *N* fragments if it has *N* + 1 restriction enzymes. Whenever such distance is consistent with that of the reference, the color of that fragment is set to yellow. When the distance on the read is smaller than that on the reference, the fra gment is in green. When the distance on the read is larger than that on the reference, the fragment is in red. The intensity of green and red represents the intensity of contraction and stretch, respectively. It is possible that the two neighboring fragments are taken as one block in the alignment, in which case their colors are the same. Left: The fragments in green in the middle of the reads (highlighted by the black box) indicate a deletion (19:40101872-40147822). But due to their not having the same boundary (some green fragments in green protrude to the right and some to the left due to two or more fragments that are aligned as one block) and the same intensity (green colors vary), the signal is weak, leading to the missing of the call by the two existing methods. Right: The fragments in orange in the middle of the reads (highlighted by the black box) indicate an insertion (1:236385430-236394679).

In this paper, we propose OMIndel, an alignment-based method combined with local assembly-like approach for indel detection. It is sensitive on calls with weak signals, an improvement over both BioNano Solve and OMSV. A test on NA12878, a healthy diploid genome, and the whole CEU trio demonstrate that OMIndel is able to detect those indels not detectable by either of the two existing methods, while simultaneously maintaining a lower or comparable false discovery rate. Furthermore, we looked into the indels that are only detectable by OM but not by sequencing-based technologies such as Illumina or PacBio, and categorized them into complex events or indels falling on repetitive regions. The method is implemented in Perl and is publicly available for download.

lignment, in which case their colors are the same. Left: The fragments in green in the middle of the reads (highlighted by the black box) indicate a deletion (19:40101872-40147822). But due to their not having the same boundary (some green sticks protrude to the right and some to the left due to the merging of two or more fragments) and the same intensity (green colors vary), the signal is weak, leading to the missing of the call by the two existing methods. Right: The fragments in orange in the middle of the reads (highlighted by the black box) indicate an insertion (1:236385430-236394679).

## 2 Methods

### 2.1 General overview of OMIndel

For aligning the reads to the reference genome, OMIndel uses the same strategy as OMSV, which integrates the results from two aligners RefAligner [24] and OMBlast [20]. From the alignment, we extract all reads that do not have high concordance with the reference (i.e., at least one of the fragment correspondences between read and reference has sizing difference larger than 2,000bp). We detect indels > 2,000 bp as this size range is the strength of OM [22], and smaller indels can be covered by other sequence technologies. The information of the discordance is recorded, including the coordinates on the reference, sizing difference, etc. The subsequent read clustering involves two steps, coarse and fine, for achieving both fast and accurate clustering. First, the coarse clustering builds a graph for the discordant records and a graph-based union-find algorithm [35] is used to find all connected components of this graph. Fine clustering is then performed on each connected component. As the coarse clustering step may have multiple indels clustered together due to false edges, for reads in one connected component, we further apply a hierarchical clustering algorithm for breaking reads into groups that are truly corresponding to the same indel. The scoring system in the hierarchical clustering is a distance ranging from 0 to 1 between each pair of the reads (0 means the two reads are exactly the same on the focal indel region, and 1 means the two are completely different). Such distance is calculated by aligning the focal region of one read to another (a dynamic programming algorithm for alignment is described below). The alignment score is normalized and subsequently taken to calculate the distance. We then classify the putative indel calls from each group into homozygous reference, homozygous variant and heterozygous variant with a variant score, followed by filtering. An outline with the cartoons is shown in Fig.2. We now turn to describing the various steps of OMIndel in detail.

**Fig. 2:**
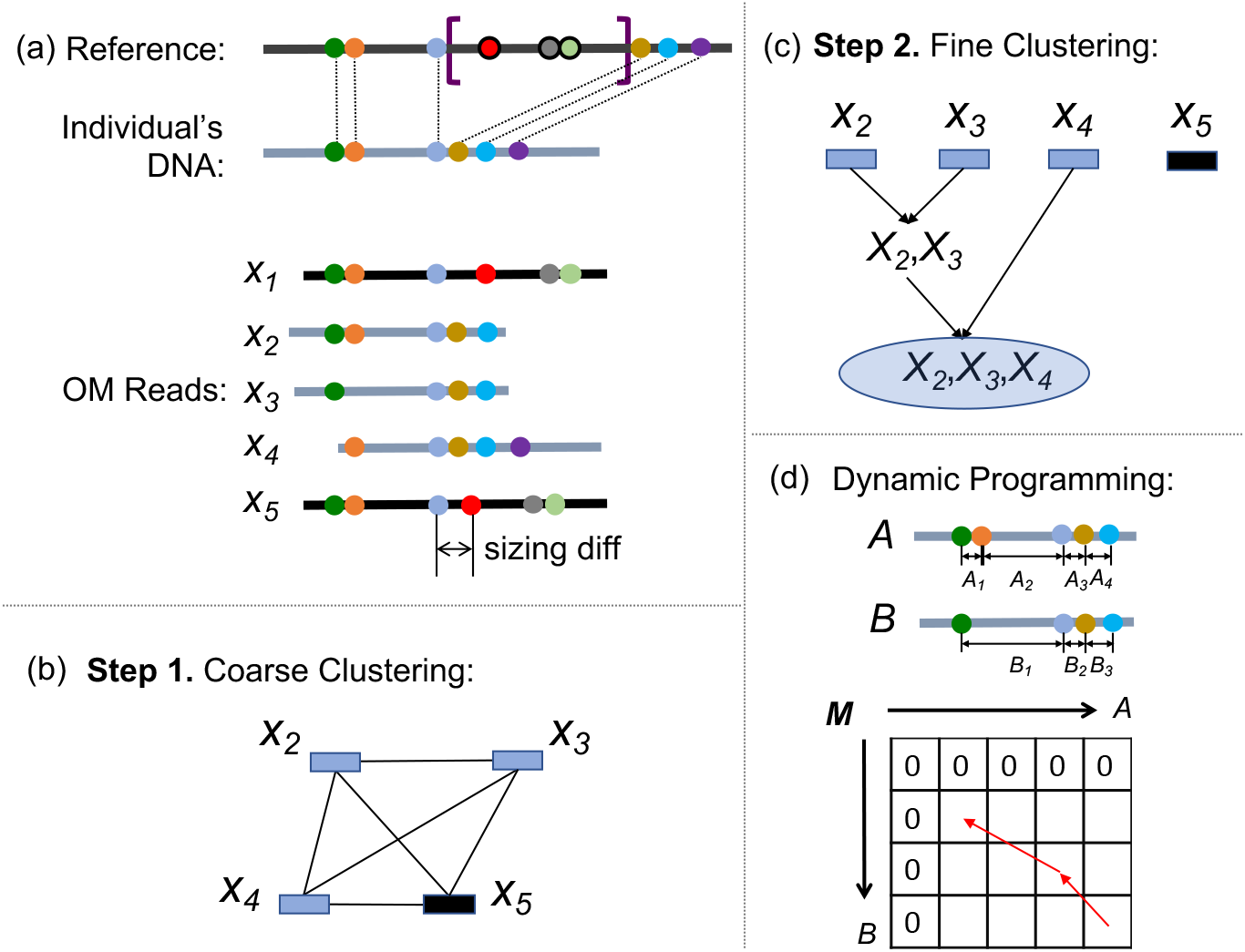
Illustration of the OMIndel method. (a) Deletion of a segment (in purple brackets) with respect to the reference genome (in black horizontal line) is shown. The three labels with black outer circles are deleted in individual’s DNA. Each label is shown in a different color for visualization. Correspondence between labels on the reference and the individual’s DNA is shown in dashed lines. Five OM reads aligned to this locus are shown: the ones in black lines (*X*_1_ and *X*_5_) come from the reference allele, and the ones in dark blue lines (*X*_2_, *X*_3_, and *X*_4_) come from a variant allele. Note that due to a sizing difference error on *X*_5_, it is selected as a variant read along with *X*_2_, *X*_3_ and *X*_4_ in coarse clustering (step 1). yet to be filtered in fine clustering (step 2). (b) Step 1. a coarse clustering is performed. *X*_2_, *X*_3_, *X*_4_, and *X*_5_ are all selected as variant reads. They represent nodes in a graph. Since they overlap with each other, they are all connected. A union-find algorithm is applied to the graph to cluster connected components, and the four reads are grouped in one cluster. (c) Step 2, a fine/hierarchical clustering is performed on individual connected components. The alignment score of each pair of two reads is calculated by a dynamic programming algorithm (described in (d)). The clustering starts from the two reads that have the smallest distance score, and stops when the distance score between two groups of reads is above a threshold. *X*_5_, due to its large distance with *X*_2_, *X*_3_, and *X*_4_, does not successfully make it into the cluster, as expected. (d) The process of dynamic programming algorithm to calculate the alignment score. *A* and *B* are two OM reads (notice that *B* does not have the orange label as that in *A* due to a missing label error). The topmost row and leftmost column of matrix *M* was initialized with zeros. The entries are filled from top left to bottom right and the la rgest value is selected as the alignment score. The traceback path (shown in red arrows) retrieves the optimal alignment. In this example, *A*_1_ and *A*_2_ are joined as a group to be compared with *B*_1_, with a gap penalty applied.

### 2.2 Union-find for coarse clustering

Before the first round of clustering, we align the OM reads to the reference (GRCh38 is used in the Results section below) in the same fashion as that of OMSV [22]. That is, the method uses integrated results from two aligners, RefAligner [24] and OMBlast [20]. We then extract the variant reads that have at least one abnormal sizing difference for all fragments in the read compared with the reference. As the two end labels on the reads have a higher error rate, sizing differences on these labels are skipped. Also, in local alignment, in case more than 5 consecutive fragments have to be aligned as one block and cannot be aligned separately, they are omitted as they probably contain large errors.

We then build an undirected graph, in which each read represents a node, and an edge links two nodes if their corresponding reads have their indicated indel coordinates overlapping with each other on the reference. A union-find algorithm [35] is applied to find all connected components in the graph, producing the clusters of reads. Following this step, each connected component is refined via fine clustering as we describe in the next section.

### 2.3 Local assembly-like approach for fine clustering

To overcome random errors in OM reads and achieve high accuracy in indel calling, one more round of clustering is performed for each connected component obtained by the previous step. Within a connected component, for a pair of reads *A* and *B*, a distance *D_AB_* is calculated, which is used in the subsequent hierarchical clustering step. The merging of clusters (in the hierarchical clustering) stops when the two clusters have their distance larger than a threshold. The distance between two groups of reads is calculated as the average distance of all pairs of reads between two groups. For a pair of reads *A* and *B*, *D_AB_* is symmetric, composed of the scores from a dynamic programming algorithm for pairwise read alignment. Specifically,

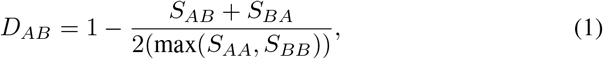

where *S_AB_* is the score of aligning read *A* to read *B* using a dynamic programming algorithm described next. When *A* and *B* are exactly the same, *D_AB_* equals zero. The maximum value of *D* is 1.

We now describe a dynamic programming algorithm for OM reads. In sequence-based pairwise alignment, dynamic programming algorithms such as Smith-Waterman [9] look for best matches between subsequences of the two sequences. A scoring system is used as a way to penalize gaps and mismatches but reward matches. In OM, the dynamic programming is designed in a similar fashion except that instead of penalizing mismatches of the nucleotide bases, we penalize the sizing difference between the two fragments. Also penalizing the indel is turned into penalizing the additional and missing labels. To allow errors that occur near each other, we take the matching of two merged sets of fragments into consideration, with a penalty to the number of fragments that are being merged. More formally, suppose OM read *A* has fragments *A*_1_,…, *A_x_* and OM read *B* has fragments *B*_1_,…, *B_y_*. For example, if OM read A consists of four coordinates (5,10,12,18), then the fragme nts are *A*_1_ = 5 (= 10 − 5), *A*_2_ = 2 (= 12 − 10), and *A*_3_ = 6 (=18 − 12); i.e., *A_i_* is the number of positions that separate the *i*-th and (*i* + 1)-th coordinates in read *A*. The following is a dynamic programming algorithm for calculating the score of optimally aligning *A*_1…*i*_ to *B*_1…*j*_ (1 ≤ *i* ≤ *x*, 1 ≤ *j* ≤ *y*), which is stored as entry *M*(*i*, *j*).

– Initialization: *M*(0, *j*) = 0 for 1 ≤ *j* ≤ *y*, and *M*(*i*, 0) = 0 for 1 ≤ *i* ≤ *x*.
– Recursion:

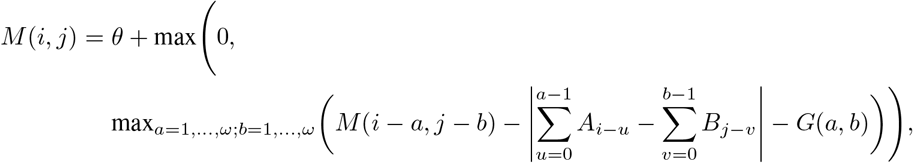 where *G* (*a*, *b*) = *σ* · (*a* + *b* − 2) is the gap penalty (*σ* is a normalizing factor that makes sizing difference penalty and gap penalty comparable), *θ* is a reward for extending the alignment to make the matrix entries positive when there is a good alignment, and *ω* (≤ min(*x*, *y*)) is a user-specific threshold on the maximum number of the fragments to be counted as one block for alignment. In the equation, the summations are taken over *u*’s and *v*’s that satisfy *i* − *u* ≥ 1 and *j* − *v* ≥ 1.
– Termination: *i* = *x* and *j* = *y*.

If two reads are on two different genomic loci, they are unlikely to have overlapping coordinates (with respect to the reference genome) on their respective OM reads and, consequently, are unlikely to belong to the same connected component as identified by the step given in Section 2.2 above. This is why the formula for *M*(*i*,*j*) does not account for the actual coordinates, but only for the “spacings” between coordinates (fragments). Finally,

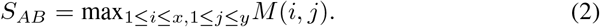

### 2.4 Genotyping

We classify each call into homozygous reference, homozygous variant and heterozygous variant by a maximum likelihood approach. The likelihood of each genotype takes both supporting read number and concordance of their indicated indel size into account. Specifically, we model the supporting read number as a Gaussian distribution (the number of reads aligned to a focal region varies and the farther that number from the mean, the smaller its frequency, hence the choice of the Gaussian distribution). We model the sizing difference of each read in a cluster as a Cauchy distribution (the sizing differences from noise have a Gaussian distribution with long tails, hence the choice of the Cauchy distribution, which is also discussed in [22]; see Fig. 3 below). The likelihoods can be expressed as follows:

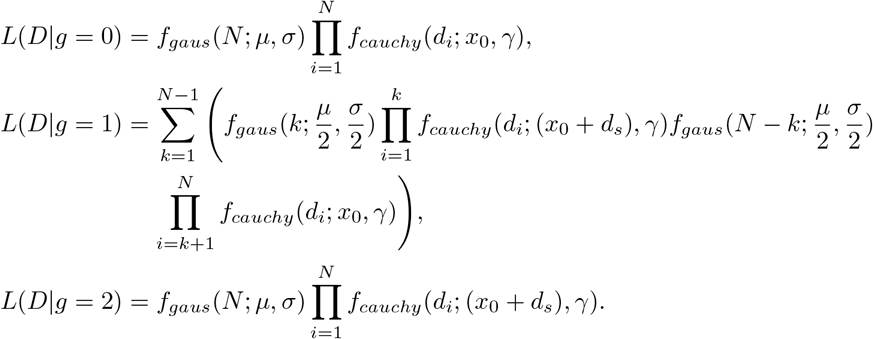

**Fig. 3:**
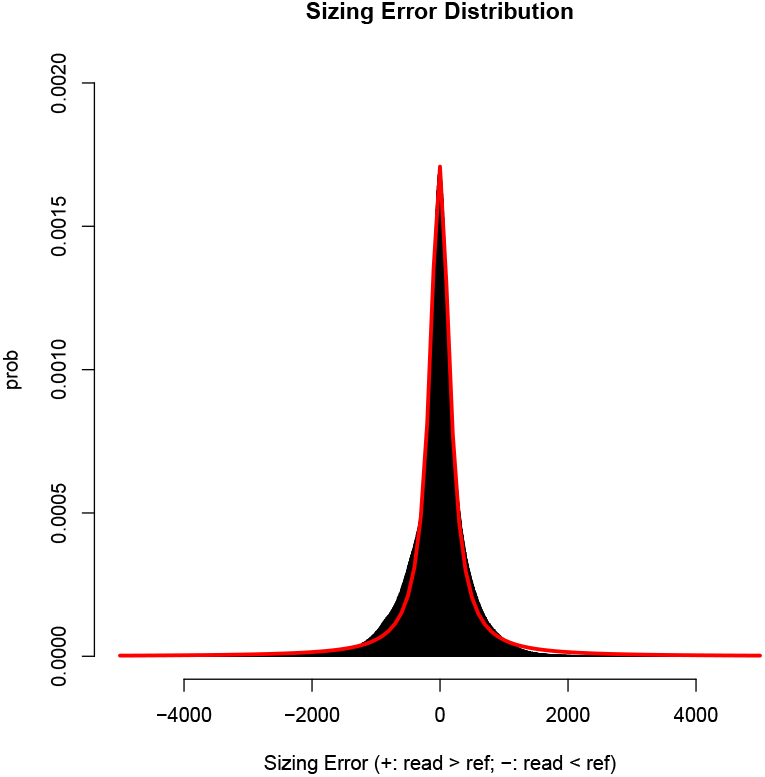
Sizing difference distribution of NA12878 as approximated by a Cauchy distribution (red curve).

In these expressions:

– *D* is the OM data (all reads aligned to the local region of interest, given in terms of their fragment length and alignment);
– *g* is the number of variant allele in the site (*g* = 0 for homozygous reference, *g* = 1 for heterozygous and *g* = 2 for homozygous variant);
– *N* is the total number of reads on the site;
– *μ* and *σ* are the parameters learned from the whole genome, representing the mean and standard deviation of the number of reads covering a site;
– *d_i_* is the inferred indel size from the *i^th^* OM read;
– *x*_0_ and *γ* are the location and scale parameters of the Cauchy distribution learned from the whole genome where the assumption is there is no indel; and,
– *d_s_* is the estimated indel size given from the previous local assembly-like step, which is the mean of the inferred indels from the reads that cluster.

For homozygous reference, there is no indel, and the location parameter of the sizing difference between read and reference is simply *x*_0_, the one learnt from the whole genome. For homozygous variant, the sizing difference of every read on the site corresponds to the same Cauchy distribution learnt from the whole genome, except that the distribution shifts to the left by *d_s_*. For heterozygous variant, suppose *k* reads support the variant and the rest of *N* − *k* reads support the reference. The variant and reference supporting read number should both be corresponding to a modified Gaussian distribution (i.e., the mean and variant are half of the *μ* and *σ*), with their sizing difference to the reference and variant Cauchy distribution, respectively.

To improve computation time, we approximate the heterozygous variant’s likelihood as follows:

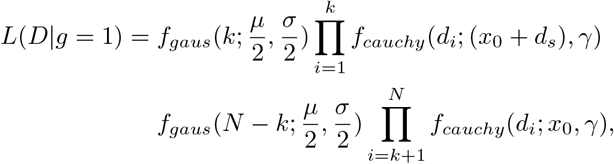

where *k* represents the number of variant-supporting reads that are clustered in the previous step, and *N* − *k* is the number of remaining reads aligned to the site. As the previous step assembles all the reads supporting the same allele, this approximation is valid as the other terms in the summation (the first equation for *L*(*D*|*g* = 1) above) are close to zero and thus can be omitted. When a read supporting the variant is wrongly clustered as a reference read, as long as its sizing difference is > 1,000bp (some weak signal exists), such omission makes a difference of only less than 6.12e-05 (through the calculation of Cauchy distribution by setting *γ* = 200). Finally, a maximum likelihood estimate of the genotype is given by

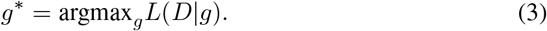

The variant score can be calculated as

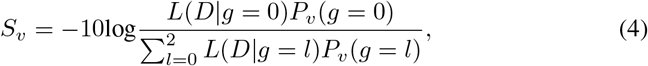

where *P_v_*(*g*) are the prior probabilities for the three genotypes, and

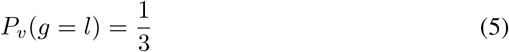

for *l* = 0,1 and 2.

## 3 Results

### 3.1 Simulated data

Our simulation process involves two steps: simulating variant alleles and simulating OM reads. In simulating variant alleles, on the in silico digested human reference chromosome 20, we simulate 50 deletions and 50 insertions. For each indel, we uniformly sample its starting label. The indel size is sampled from a Cauchy distribution (locality = 0, scale = 300) and is at least 2,000bp. We use a Cauchy distribution to simulate the real situation where medium-sized indels outnumber large indels. Labels that are covered in the deleted area are also deleted. For insertions, we simulate the inserted labels such that the distance between the current and the next inserted label is drawn from a Poisson distribution, where the mean is the average distance between two labels in the real case (10kbp). This process of simulating the inserted label is repeated until no more labels can be sampled from the simulated insertion size. To avoid sampling overlapping indels, we constrain the distance between each pair of neighboring indels to be >100kbp.

In simulating OM reads, we learned the statistics, including read length, error rates and sizing difference from the real data (CEU trio) and approximated with distributions described below. We simulated three total coverages: 80x, 100x, and 120x and four variant allele fractions (VAFs): 0.2, 0.3, 0.4 and 0.5, resulting in 12 genomes, each having one variant allele and one reference allele. In simulating a read from a given allele (reference or variant), we uniformly sample the starting point. From the real data, we estimated the median of read’s length to be about 200kbp. Since the minimum read length starting to contribute to SV detection is 150kbp [3], we set the length of the read to be *l*_0_ + *l_r_*, where *l*_0_ is 150kbp and *l_r_* is sampled from a Poisson distribution with mean at 50kbp. Next, we learned the error profiles from the high-confidence alignments (alignments whose reads have ≥ 12 labels and clipped end is ≤ 4 labels). The following items are the statistics learned for the three error types.

– Missing label error rate’s median is 0.05 (similar to that reported in [22]);
– Additional label error rate is one per 200kbp;
– Sizing difference’s distribution is Cauchy-like as it has long tails (Fig. 3). The Cauchy distribution’s parameters are approximated to be locality = 0 and scale = 200 (this is similar to [22]).

We iterate this process until we simulate enough reads for the desired depth at this allele for a specific VAF and total coverage.

We applied OMSV, BioNano Solve and OMIndel to the simulated data, and measured the recall and precision of the three methods. To reduce the effect of randomness, for each total coverage and VAF, we simulated five data sets of OM reads, in order to obtain a set of accurate measurements. Fig. 4 and Fig. 5 show the recall and precision, respectively, of all three methods. For all coverages, OMIndel has higher recall than OMSV at all VAFs while maintaining comparable precision. Similarly, OMIndel is advantageous over BioNano Solve on almost all VAFs for both deletion and insertion on both recall and precision. BioNano Solve has a slight advantage over OMIndel for insertion when VAF is at 0.5 for 100x, or when VAF is at 0.2 or 0.3 for 80x, at the cost of much lower precision. We observe that at low VAFs, with the increasing of the coverage, BioNano Solve’s recall decreases. This shows the instability of BioNano Solve when reference reads greatly outnumber variant reads. Overall, our algorithm has the advantages on recall at small VAFs, with comparable precision with the other two methods for higher VAFs.

**Fig. 4:**
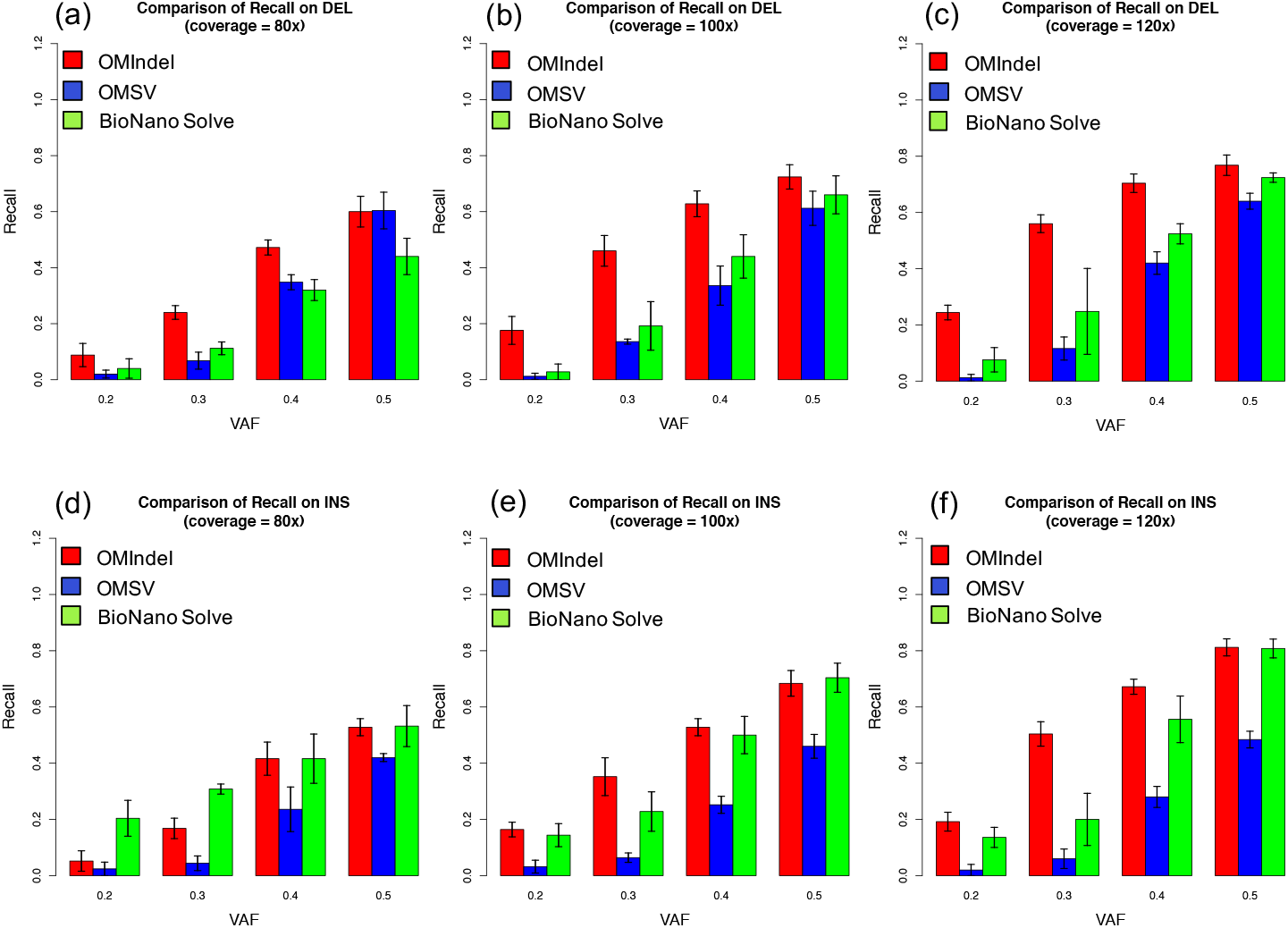
Comparison of recall on simulation data on VAF = 0.2, 0.3, 0.4, 0.5 among OMIndel, OMSV and BioNano Solve, for deletion with coverage at (a) 80x (b) 100x (c) 120x and insertion with coverage at (d) 80x (e) 100x and (f) 120x. Height of the bars represents the mean; the error bars represent the range within one standard deviation whenever it is within [0,1].

**Fig. 5:**
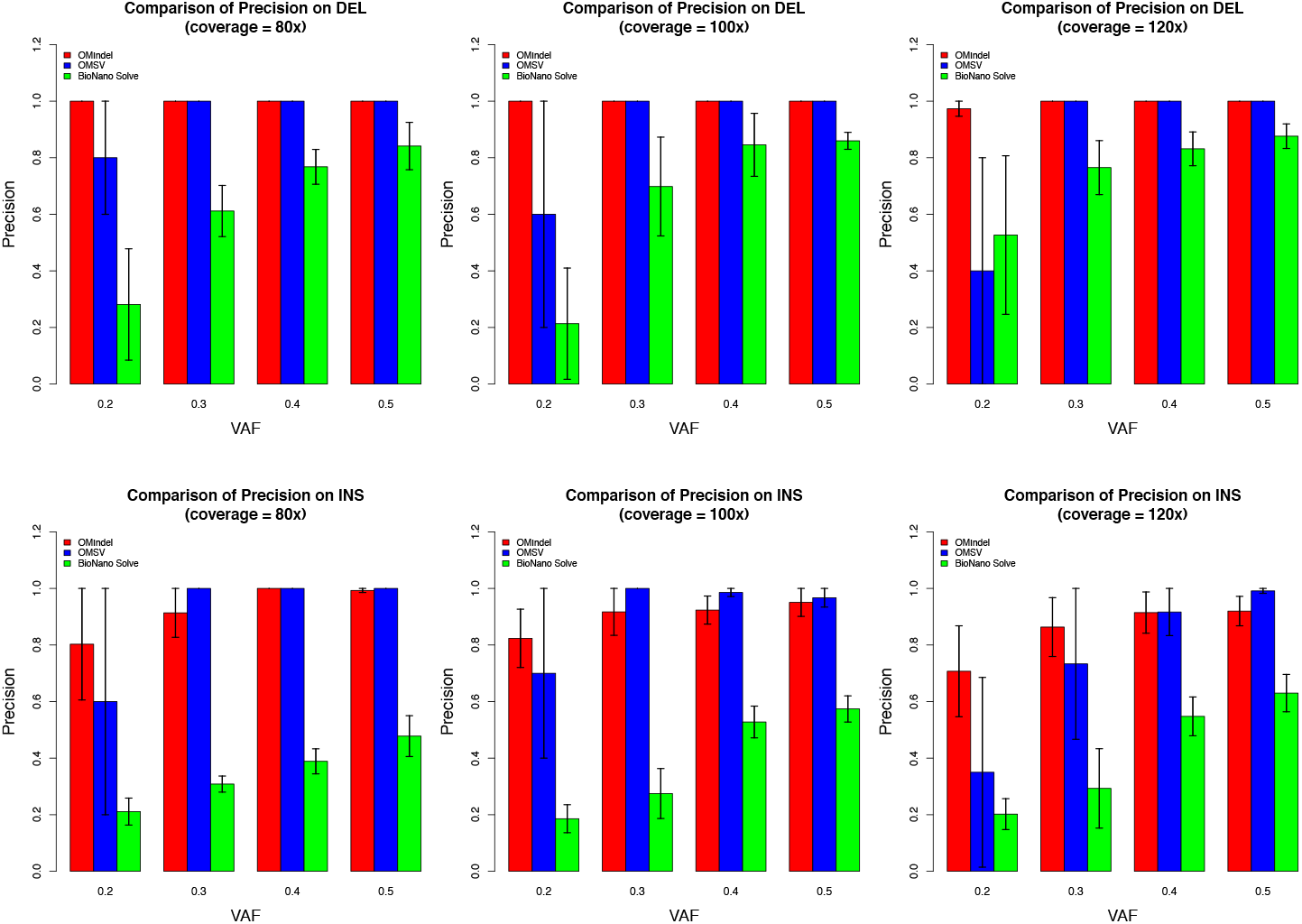
Comparison of precision on simulation data on VAF = 0.2, 0.3, 0.4, 0.5 among OMIndel, OMSV and BioNano Solve, for deletion with coverage at (a) 80x (b) 100x (c) 120x and insertion with coverage at (d) 80x (e) 100x and (f) 120x. Height of the bars represents the mean; the error bars represent the range within one standard deviation whenever it is within [0, 1].

It is important to note here that BioNano Solve does not report the indel sizes along with the predictions, which is the reason why we cannot report the recall and precision of the methods broken down by indel sizes (as the indel size could often be a factor in a method’s performance).

We investigated whether the low recall of OMSV is due to false alignments. We generated a “ground truth” alignment file given our knowledge of the indel and read errors for one of the twelve cases (VAF=0.5, total coverage=120x). We then applied OMSV to the true alignments and found that while maintaining a high precision (1 for both deletion and insertion), the recall for deletion and insertion are respectively 1 and 0.98, compared with 0.64 and 0.44 from the alignment that has errors. This shows that OMSV’s recall was greatly affected by erroneous alignments.

Lastly, we evaluated the computational cost for the three methods (Table 1) for one of the simulation data sets (coverage=100x). OMSV is the fastest while requiring the lowest amount of memory. OMIndel is the second (around 30 times faster than BioNano Solve while requiring relatively small amount of memory). BioNano Solve requires a large amount of CPU hours as well as memory. Here the CPU hours are the total ones if parallelization is applied for a fair comparison, and the same to memory.

**Table 1:** Comparison of computational cost on simulation data.

|  | OMIndel | OMSV | BioNano Solve |
| --- | --- | --- | --- |
| CPU hours | 5 hours | 0.5 hours | 140 hours |
| Memory | 4G | 1G | 20G |

### 3.2 Empirical data

We applied OMIndel to NA12878 (VAF of 0.5 for heterozygous events, and ~90x coverage), and called 479 deletions and 700 insertions. Comparing the calls with those of OMSV and BioNano Solve, OMIndel uniquely called 62 (13%) deletions and 87 (12%) insertions (Venn diagram in Fig. 6a and b), in which 37 (60%) deletions and 77 (89%) insertions are also called by either parents (NA12891 and NA12892) or overlap with orthogonal sequencing-based calls. We construct the orthogonal sequence-based calls such that it is a deduplicated union set of indels from Delly [32], PacBio calls generated in [30], 10X calls (ftp://ftp-trace.ncbi.nlm.nih.gov/giab/ftp/data/NA12878) and hybrid methods including HySA [10], svclassify [29] and metaSV [26]. We found here that only 25 (5.2%) deletions and 10 (1.4%) insertions can be validated neither by the parents’ calls nor by orthogonal sequence-based calls. The estimated precision of OMIndel is therefore 94.8% for deletion and 98.6% for insertion, compared with 93.4% and 99.7% for OMSV and 93.0% and 95.4% for BioNano Solve. It was observed that among those numbers, only OMSV’s insertion detection has around 1% advantage of precision over OMIndel. However, it is at the cost of missing 127 (22.8%) validated insertions shared by both OMIndel and BioNanoSolve. In addition, Fig. 6e and f show that OMIndel can potenti ally complement or even outperfoπn OMSV and BioNano Solve in teπns of detecting novel calls validated by sequence-based method (13 for deletion and 26 for insertion, compared with 13 and 11 for OMSV, and 53 and 62 for BioNano Solve).

**Fig. 6:**
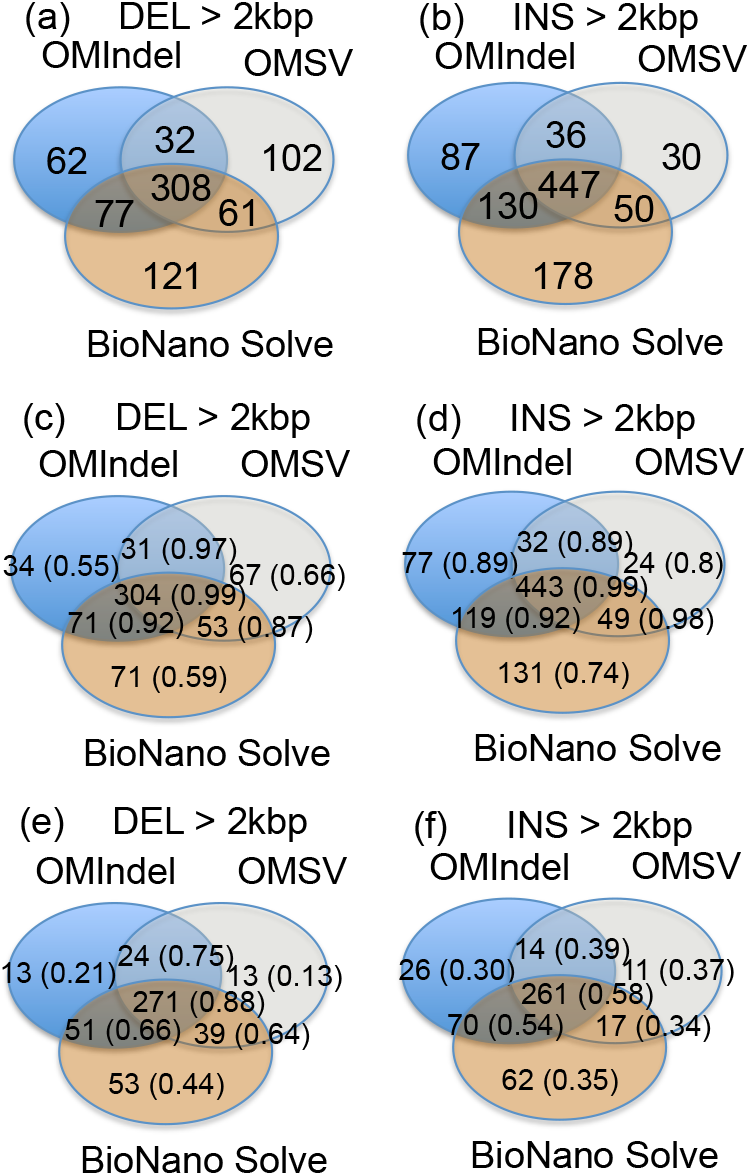
Venn diagram comparing the indels of OMIndel, OMSV and BioNano Solve. (a) and (b) show the numbers of deletion and insertion in the Venn diagram. (c) and (d) show the number of calls that can be validated by the parents’ OM calls for deletion and insertion, respectively. (e) and (f) show the number of calls that can be validated by the orthogonal sequenced-based method for deletion and insertion, respectively. The corresponding percentages over the total call are in parenthesis.

We further looked into OMIndel unique and validated calls (named set A), and compared with the number of supporting reads between those are shared (set B). We found set A has a much smaller number of variant supporting read number (Table 2) than that of set B. Along with the recalls from simulation, OMIndel has been proven to be advantageous in calling indels with weak signals.

**Table 2:**
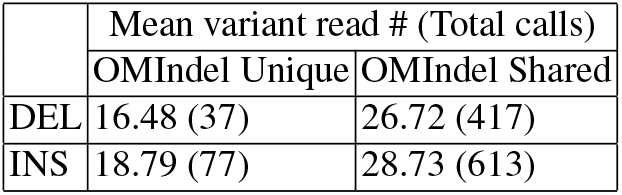
Comparison of mean variant supporting read number between unique validated calls and shared calls. In the parenthesis are the total call number within the category.

The CEU trio data provided us the opportunity to evaluate our genotype accuracy compared with the other two methods. Table 3 listed the accuracy of proband’s genotype given parents’ calls and corresponding genotypes according to the mendelian inheritance rule. Except on deletion, when OMIndel’s genotype accuracy ties with that of BioNano Solve, OMIndel outperforms the two methods on both insertion and deletion. Particularly, we observed that OMSV’s genotype accuracy is pretty low, as compared with its high precision in making a call. This particularly demonstrates OMSV’s disadvantages in extracting supporting variant and reference reads corresponding to a variant call when mis-alignments are involved.

**Table 3:** Comparison of genotype accuracy.

|  | GT Accuracy |  |
| --- | --- | --- |
|  | DEL | INS |
| OMIndel | 0.75 | 0.83 |
| OMSV | 0.63 | 0.73 |
| BioNano Solve | 0.75 | 0.77 |

### 3.3 Investigating novel calls missed by sequence-based methods

We further compared OMIndel calls with the sequence-based calls described above for both deletions and insertions (Fig. 7). We investigated the novel calls missed by all sequence-based methods but can be found in parents’ calls. Of the 479 deletions, 120 (25%) are missed by sequence-based methods, of which 86 (72%) are validated by parents’ calls. Of the 700 insertions, 329 (47%) are missed by sequence-based methods, of which 296 (90%) are validated by parents’ calls. We randomly selected 20 deletions and 20 insertions that are novel to sequence-based calls but called in parents. We found these novel indels are missed by sequencing-based methods mainly because they fall into repetitive regions (Fig. 8) or they are complex events (Fig. 9). In all, these events are missed by Illumina or PacBio based methods mainly because the variant signals are very weak.

**Fig. 7:**
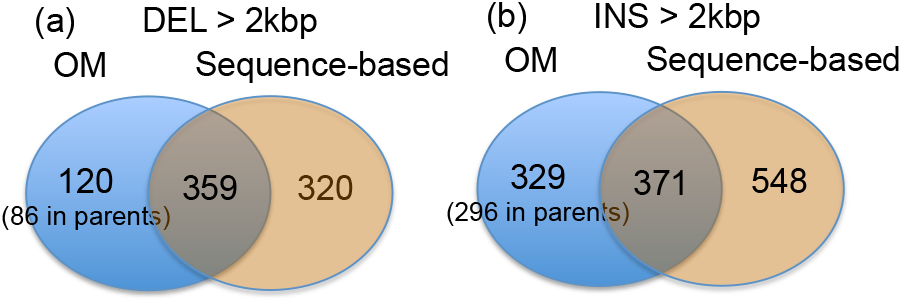
Venn diagram comparing sequence-based indels and OMIndel calls on OM. The numbers in parenthesis are those that are also called by parents.

**Fig. 8:**
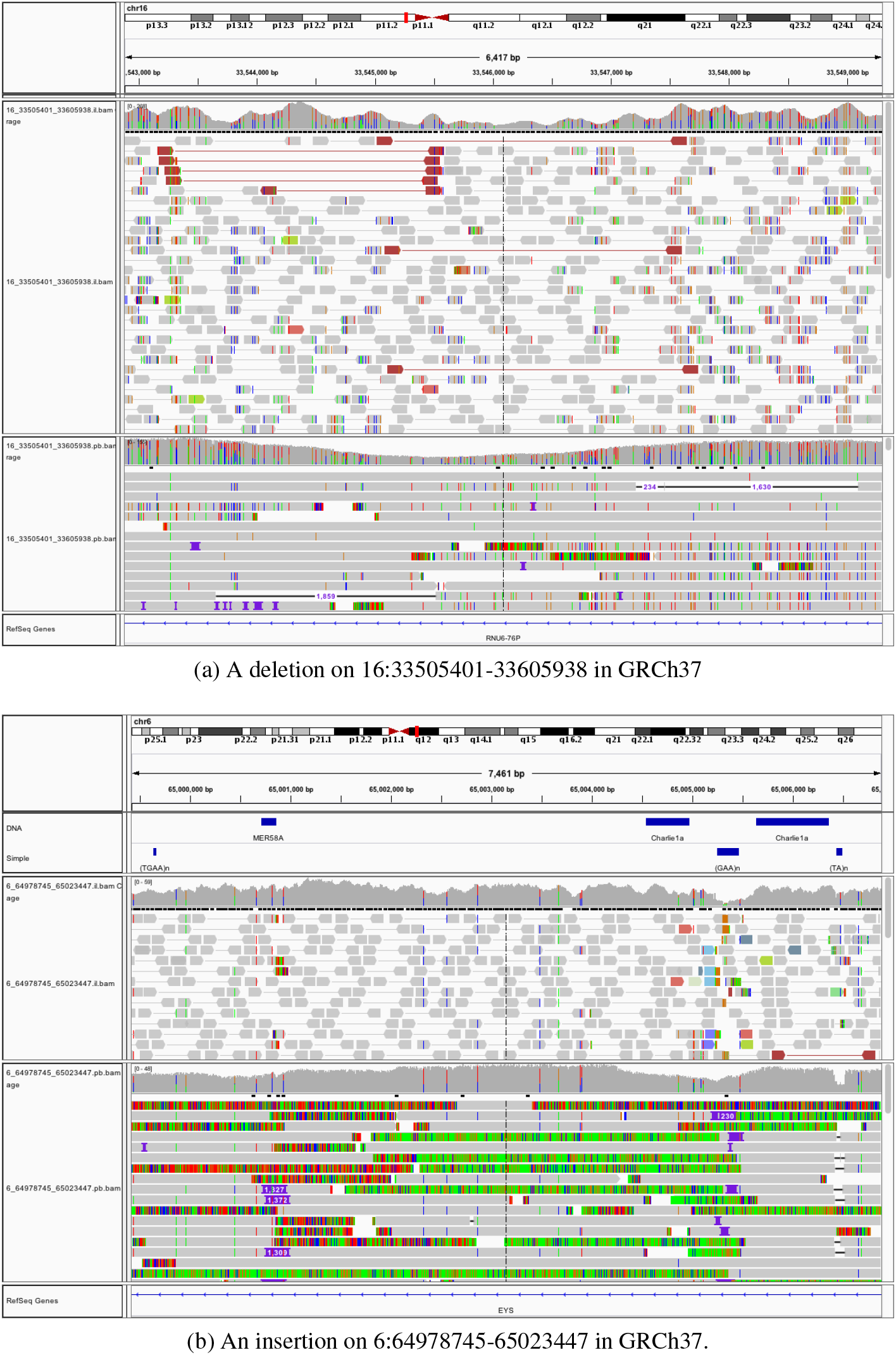
IGV of complex indels that are also called by parents but missed by sequence-based methods. In all IGVs, the upper and bottom panels show the alignment of Illumina and PacBio reads, respectively.

**Fig. 9:**
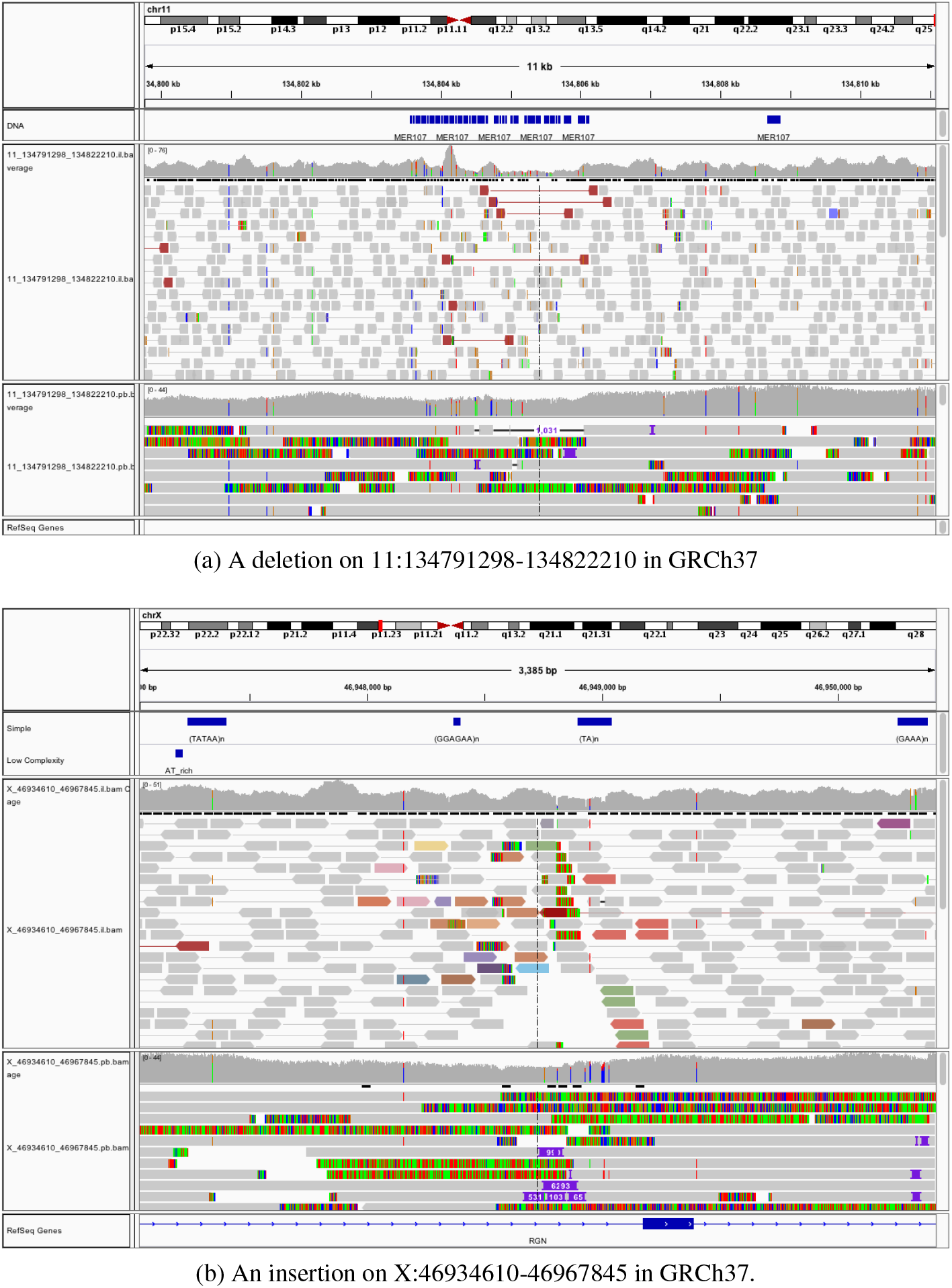
IGV of indels overlapping with repetitive regions. These indels are also called by parents but missed by sequence-based methods.

## 4 Discussion

With the long reads provided by OM data, it is of most interest to know what OM can deliver as compared with other technologies. While this study is focused on answering this question, we found the existing tools on OM are limited in handling the indels with weak signals. Our proposed OM-based targeted assembly approach, OMIndel, falls in the same paradigm of that in TIGRA [5] and HySA [10].

In simulations, we found that BioNano Solve is only advantageous over OMIndel on high VAFs and insertions when measuring recall. This is consistent with our prior assumption that a *de novo* assembly-based approach is better at detecting large insertions than alignment-based or local assembly-based approach when there are enough reads representing the variant. The recall corresponding to different insertion sizes is yet to be summarized for alignment-based or assembly-based approaches.

We summarized some major categories where OM has its unique advantage of detecting indels over sequencing-based technologies. However, further investigation is needed so that such list of categories can be comprehensive as a reference for sequencing. We also observed that there are quite a few sequence-based calls (47.1% for deletions and 59.6% for insertions) missed by our OM-based method. Further investigation will facilitate exploring why OM missed the indels called by sequence-based methods. This may further help to increase the recall of OM-based algorithms. It is also valuable to identify OM’s limitations, i.e., which indels are beyond OM’s detection ability. This could be done by looking into the calls unique to sequence-based methods.

We acknowledge that the error profiles unique to OM data have not been fully investigated in this study. As discussed by [22], missing and additional labels are indicative of small indels, whereas sizing differences are indicative of large indels. In this study, we focused only on large indels (indels > 2000bp) by modeling sizing difference distribution in our genotyping algorithm. Detecting smaller indels requires further investigation where the missing and additional label errors need to be modeled.

Finally, this paper’s scope is limited only to indel detection. OM’s long reads have advantages in detecting inversions, translocations, or complex events such as chromothripsis and chromoplexy. With all of these explored, a comprehensive characterization of the human genome could potentially be achieved.

## 5 Conclusions

We proposed OMIndel, a method that utilizes OM reads to detect large indels. It differs from the previous two methods in that it is alignment-based but follows a local assembly-like fashion, so that it can simultaneously detect indels with weak signal as well as maintain a low FDR. We applied OMIndel to both simulated and real data, and found that it is advantageous over the other two OM-based methods, OMSV and BioNano Solve, by detecting those indels that have weak signals while maintaining a higher or comparable precision. We also manually inspected the indels unique to OM but missed by sequence-based methods. We found that they fall into either a category of complex events or at repetitive regions. OMIndel is freely downloadable online, and we expect that with the increasing availability of samples having OM data and the decreasing cost of OM technology, this tool can be widely used for SV detection.

## Acknowledgements

We thank Drs. Ken Chen at MD Anderson Cancer Center and Feng Yue at Pennsylvania State University for discussions and data sharing. We would like to thank the Human Genome Structural Variation Consortium. Special thanks to Dr. Charles Lee, Dr. Evan Eichler, Dr. Jan Korbel and Dr. Mark Chaisson for their comments on OM analysis. We used the Extreme Science and Engineering Discovery Environment (XSEDE), which is supported by National Science Foundation grant number ACI-1548562.

## Footnotes

* This work was supported in part by the National Cancer Institute (NCI) grant R01-CA172652 and National Human Genome Research Institute (NHGRI) grant U41-HG007497-01 to Ken Chen at MD Anderson Cancer Center, and the National Cancer Institute Cancer Center Support Grant P30-CA016672 to the MD Anderson cancer center.

